# IntAct-U-ExM: Ultrastructure Expansion microscopy of actin networks via an internally-tagged actin

**DOI:** 10.1101/2025.05.14.654030

**Authors:** Anubhav Dhar, Nishaant Kumar Palani Balaji, Sanjana Mullick, Sucheta Dey, Angana Ghosh, Deepak Nair, Sudarshan Gadadhar, Saravanan Palani

## Abstract

Expansion microscopy (ExM) has revolutionized super-resolution imaging in cell biology due to its simple and inexpensive workflow. The use of ExM has revealed several novel insights into the nanoscale architectures of cellular protein complexes, especially the microtubule cytoskeleton in model and non-model systems. Despite tremendous progress in expansion microscopy protocols that preserve cellular ultrastructure (U-ExM), compatible probes for imaging actin isoforms with U-ExM are still lacking and have hindered the study of diverse actin isoforms and networks across model systems. Here, we use IntAct, an internally tagged actin that incorporates into cellular actin networks, to develop and optimize U-ExM of diverse actin network types in both yeast and mammalian cells. Using expression of ALFA-tagged IntAct variants in yeast and mammalian cells, we show robust visualization of actin patches, cables, and rings in yeast and diverse actin networks such as actin cortex, stress fibers, filopodia, lamellipodium in mammalian cells at improved resolution. We also detect transient nuclear actin filaments using IntAct-U-ExM underscoring the advantages offered by our approach to image understudied actin structures. Overall, we demonstrate the effectiveness of IntAct-U-ExM for performing super-resolution imaging of various actin structures in an isoform-specific manner and highlight the potential of IntAct to study the nanoscale organization of diverse actin cytoskeletal networks across species.

**Graphical Abstract:** 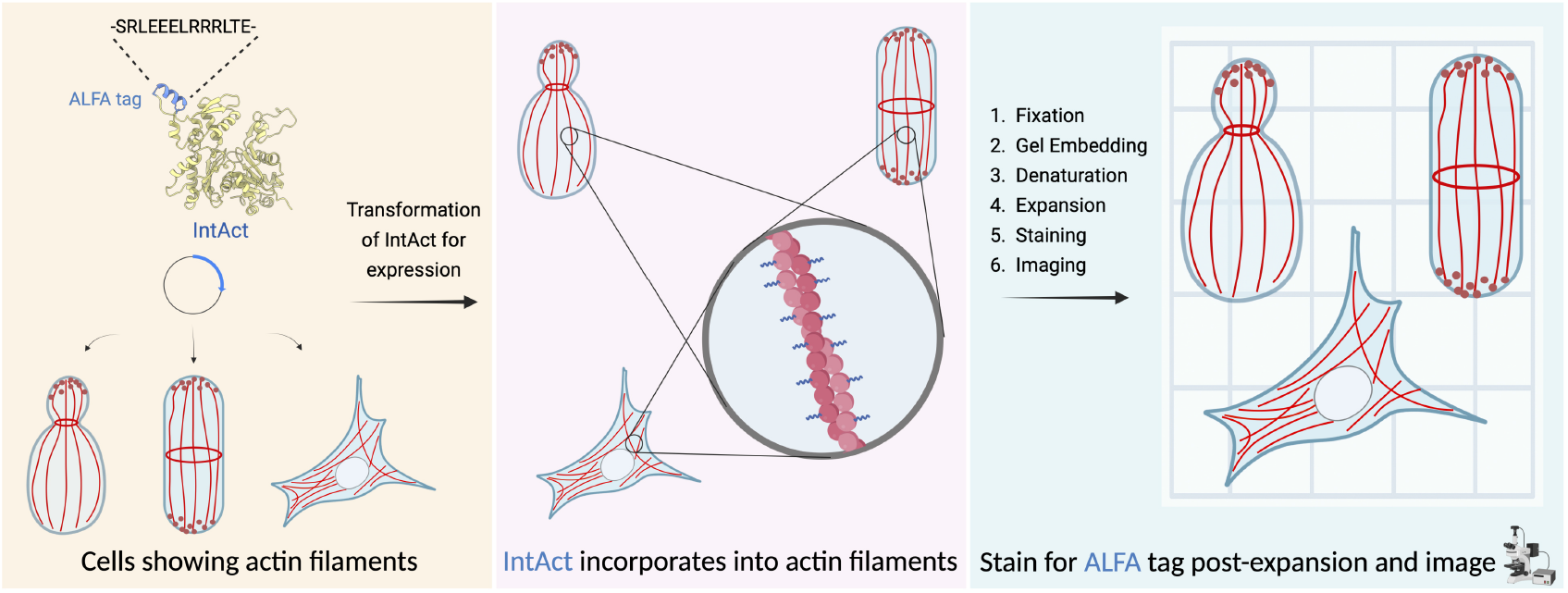

## Introduction

Super-resolution fluorescence microscopy techniques have revolutionized the field of cell biology over the last two decades^1,2^. The need to visualize biomolecules at ever increasing resolution below the diffraction limit has led to the development of advanced techniques like STED^3,4^, SIM^5–7^, STORM^8^, PALM^9,10^, DNA-PAINT^11–13^, MINFLUX^14,15^. A recent addition to this list, Expansion microscopy, began as a qualitative method for imaging biological samples in the Boyden lab at MIT in 2015^16^, but has evolved rapidly in the last decade to offer spatial information comparable to super-resolution microscopy. It is a first-of-its kind technique which uses physical expansion of the biological specimen instead of advanced optics and computation to circumvent the diffraction limit^17,18^. The Guichard and Hamel lab at University of Geneva modified this protocol in 2019, where they enabled preservation of the ultrastructure of cellular organelles during the expansion procedure that enabled visualization of diverse cellular ultrastructures at nanometer resolution (U-ExM), which has since become one of the most routinely used protocol for expansion microscopy^19^. Over the years, several iterations of the original ExM protocol have significantly made the workflow more user-friendly, accessible to any molecular biology laboratory, improved reproducibility, and extended applicability for various cell types and species^20–24^. Innovations in the chemistry underlying expansion has enabled advancement in expansion factors and combination with other super-resolution modalities have greatly increased the achievable molecular resolution^19,22,23,25–29^.

Expansion microscopy involves embedding and crosslinking the biological specimen in a swellable hydrogel that physically expands in volume by absorbing water. The sample is denatured after embedding in the gel to allow isotropic expansion of the biological specimen with minimal changes to the cellular architecture. The biomolecule-of-interest can be visualized by staining the hydrogel with antibodies or other stains pre/post-expansion of the sample and imaging the sample with any conventional widefield or confocal microscopes. Denaturation and expansion can make epitopes more accessible, allowing many antibodies, especially those that recognize denatured proteins used in western blots to work well in expansion microscopy. Thus, there exists a net-positive trade-off where some antibodies or staining reagents may lose binding capability due to a denatured epitope but overall, the process tends to improve staining compatibility. These advances have allowed visualizing various cellular structures including the cytoskeletal filaments of tubulin across species^30,31^. Despite this progress, visualizing actin via expansion microscopy has remained challenging due to a lack of probes. Actin cytoskeleton plays diverse roles in countless cellular processes and the nanoscale organization of actin and actin-binding proteins within different actin networks is an active area of research^32–40^. Recent studies have developed modified phalloidin conjugates that enable visualization of F-actin structures in expanded samples^41,42^. However, due to their incompatibility with heat denaturation and post-expansion labelling in the U-ExM protocol, these probes cannot be used in U-ExM workflows and do not provide isoform-specific information on actin. Apart from modified phalloidin probes, anti-actin antibodies can be used to label actin post-expansion^43,44^, but they suffer from cytoplasmic background, poor labeling post-denaturation of epitope and high-linkage error. In addition, anti-actin antibodies require careful optimization of fixation and labeling conditions^43–45^, and unlike tubulin, pan anti-actin antibodies that can reliably label actin post-expansion across species have not yet been reported^30^, making them a less ideal candidate. Thus, there is an urgent need to develop universal and versatile probes for actin compatible with U-ExM.

In this study, we have developed and optimized a protocol for performing U-ExM of actin isoforms in yeast and mammalian cells. Previously, we have reported a permissive site for epitope tag insertion within the actin protein (T229/A230), called “IntAct”, which shows isoform-specific incorporation into native actin filaments across species^45^. Here, by expressing IntAct actin variants with an ALFA tag^46^, we achieve clear post-expansion labelling of specific actin isoforms in mammalian and yeast cells using the nanobody against the ALFA tag^46^ (NbALFA) with a good signal-to-noise ratio. Our IntAct strategy enables visualization of actin network organization below the diffraction limit in cells from diverse model systems in an isoform-specific manner, opening vast possibilities towards understanding the nanoscale architecture and functions of actin filaments across various network types.

## Results and Discussion

### IntAct enables Ultrastructure Expansion microscopy of actin networks in yeast cells

The yeast actin cytoskeleton consists of three major actin structures: 1) Actin cables - bundles of linear actin filaments nucleated by formin proteins^47,48^, 2) Actin patches - branched actin networks at endocytic sites nucleated by the Arp2/3 complex^49,50^, 3) Actin rings - bundled actin filaments at the mother-bud neck nucleated by formins^51,52^. Previously, we have shown that IntAct can incorporate in all these three actin structures when expressed in budding and fission yeast^45^. We reasoned that IntAct’s efficient incorporation into native actin structures, minimal disruption to filament dynamics, and the known stability and versatility of the small internal ALFA tag to perform various applications could allow reliable staining with NbALFA^46,53^, enabling super-resolution analysis of actin isoforms using expansion microscopy. To test this, we used *S. cerevisiae* and *S. pombe* strains that expressed their native IntAct proteins from an exogenous plasmid copy and performed U-ExM using a recently optimized protocol for yeast^54,55^. Imaging *S. cerevisiae (S.c.)* and *S. pombe* (*S.p*.) cells revealed staining of actin structures with NbALFA-Alexa647 in both the non-expanded and expanded cells. Interestingly, detection of actin cables was significantly better in expanded samples, suggesting increased accessibility of the ALFA tag for NbALFA binding post-denaturation and expansion **(Fig. 1A, 1B)**. In contrast, Alexa488-Phalloidin could stain actin structures only in non-expanded cells and no clear staining was observed in expanded cells **(Fig. 1A, 1B)**, consistent with the known incompatibility of fluorescent-conjugated phalloidin dyes with expansion microscopy^42^. We measured cell area and dimensions and observed an average expansion factor of 4.92 for *S. cerevisiae* **(Fig. 1C)** and 4.58 for *S. pombe* **(Fig. 1D)**, as compared to non-expanded cells.

**Figure 1.**
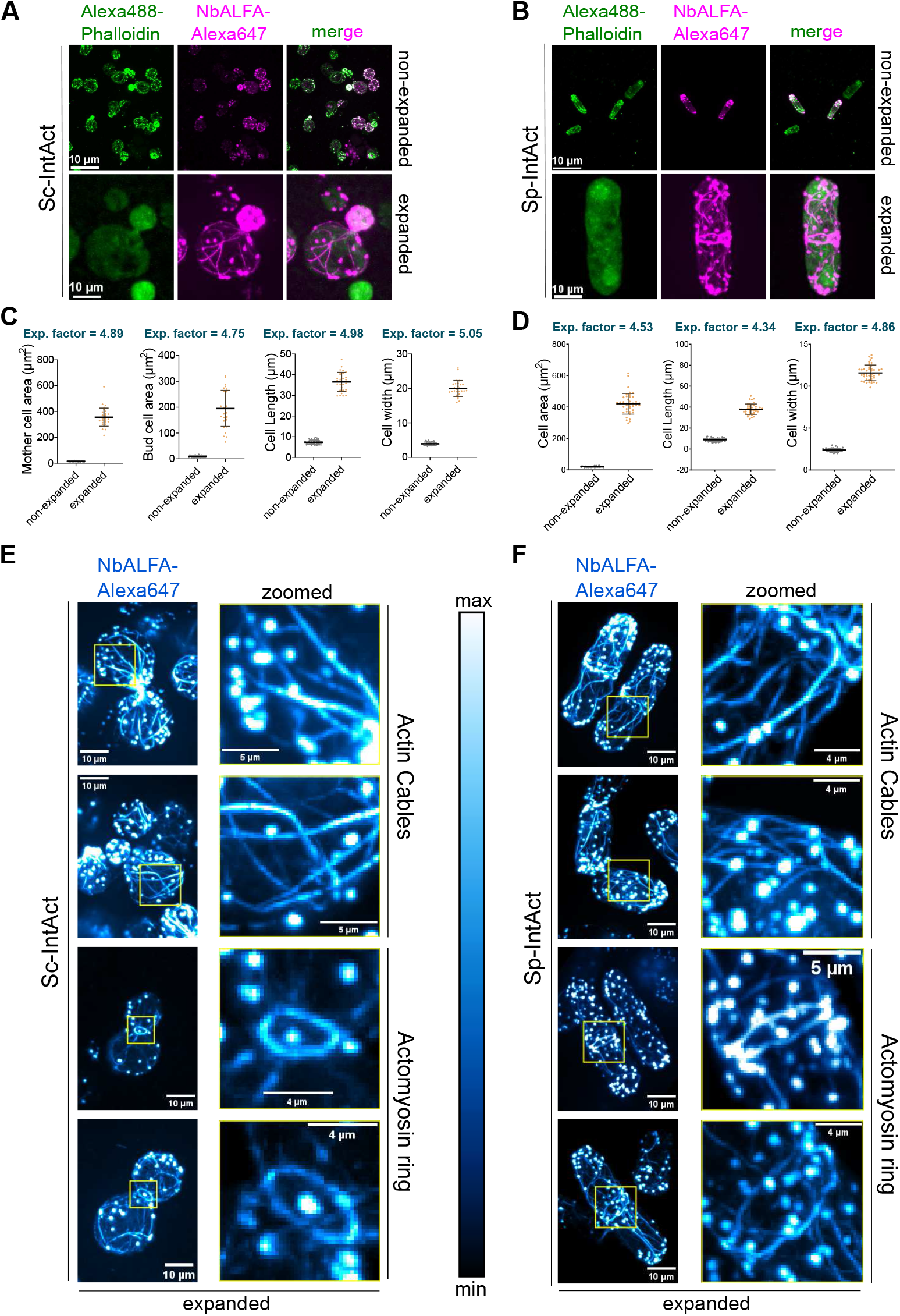
IntAct-U-ExM enables visualization of actin patches, cables and rings in budding and fission yeasts. **(A)** Representative maximum intensity projected images of non-expanded and expanded *S. cerevisiae* cells expressing Sc-IntAct stained as indicated. **(B)** Representative maximum intensity projected images of non-expanded and expanded *S. pombe* cells expressing Sp-IntAct stained as indicated. **(C)** Plots representing measurements of *S. cerevisiae* mother cell area, bud cell area, cell length (mother tip to bud tip), mother cell width in non-expanded and expanded samples and calculated expansion factors. **(D)** Plots representing measurements of *S. pombe* cell area, cell length, cell width (at cell equator) in non-expanded and expanded samples and calculated expansion factors. **(E)** Representative maximum intensity projected images of expanded *S. cerevisiae* cells stained with NbALFA-Alexa647 showing actin patches, cables, and rings. **(F)** Representative maximum intensity projected images of expanded *S. pombe* cells stained with NbALFA-Alexa647 showing actin patches, cables, and rings. (LUT display range is indicated as a vertical bar in the figure)

Despite successful staining of actin structures post-expansion, we observed that actin cables in both yeasts were not well preserved and showed discontinuous staining along their length **(Fig. S1A, S1B)**. To improve this, we compared NbALFA staining in cells fixed with 4% formaldehyde (FA) or a mix of 4% formaldehyde (FA) + 0.1% glutaraldehyde (GA), which has been shown to improve preservation of native microtubules during U-ExM previously^43^. We observed that cells fixed with 4% FA + 0.1% GA showed significantly better preservation and uniform-staining of actin cables **(Fig. S1A, S1B)**. With this optimized protocol, we successfully and consistently imaged actin patches, actin cables, and actomyosin rings in the yeasts *S. cerevisiae* **(Fig. 1E)** and *S. pombe* **(Fig. 1F)**. Since cytoplasmic actin cables are the most difficult to detect actin structures in yeast due to lower actin density as compared to actin patches, we treated *S. pombe* cells with the Arp2/3 inhibitor, CK666, which results in loss of actin patches and increase in number and intensity of actin cables^56^. The detection of actin cables was significantly better upon CK666 treatment **(Fig. S1C)** and could be used to achieve a clear organization of actin cables, demonstrating the robust use of IntAct for imaging actin networks subject to chemical perturbations. The actin cables post-expansion showed an average scaled FWHM of 74.89 ± 12.15 nm for *S.c*. **(Fig. S1D)** and 66.17 ± 9.08 nm for *S.p*. **(Fig. S1E)**. The cytokinetic actin rings showed a diameter of 4.15 *µ*m ± 0.56 *µ*m for *S.c*. **(Fig. S1F)** and 10.80 *µ*m ± 0.61 *µ*m for *S.p*. **(Fig. S1F)**, corroborating a ∼4.0-4.5 expansion factor. Overall, the above results demonstrate the application of IntAct to enable U-ExM of actin structures in yeast and its immense potential for quantitative super-resolution imaging of actin networks across the fungal kingdom.

### IntAct enables isoform-specific expansion microscopy of actin networks in mammalian cells

The success of IntAct in enabling U-ExM of yeast prompted us to test its applicability in cultured mammalian cells which harbor 6 isoforms of actin that express in different tissue types^32,57^. We used human osteosarcoma U2OS cells and specifically expressed the human non-muscle beta (β)- or gamma (γ)-IntAct isoforms in these cells from a transiently transfected exogenous plasmid. Transfected cells were prepared for U-ExM using a previously described protocol for human cells^19^ and imaged with either an epifluorescence or a spinning-disk confocal microscope. We observed clear staining of actin filaments in expanded U2OS cells expressing either β-IntAct and γ-IntAct **(Fig. 2A, 2B)**. Consistent with previous studies^42^ and our experiments with yeast, phalloidin only stained actin structures in non-expanded U2OS cells and did not stain actin structures post-expansion in U2OS cells **(Fig. 2A, 2B)**. The cells showed isotropic expansion with an average expansion factor of 3.29 and 3.84 for β- and γ-IntAct expressing U2Os cells as measured by the increase in nucleus area **(Fig. 2C, 2D)**. The lower expansion factor observed for nuclei is consistent with recent studies which suggest that all cellular compartments don’t expand with the same factor and changes in cross-linker composition can help mitigate such effects^58^. We consistently observed robust staining of β- and γ-IntAct filaments with clearly improved resolution in various actin networks such as stress fibers^59^, actin cortex^60^, lamellipodium^61^, filopodia^62^, etc. throughout the volume of expanded U2OS cells **(Fig. 2E, 2F, S2A)**. To validate these results in another mammalian model, we expressed and imaged human beta (β)- or gamma (γ)-IntAct isoforms in non-expanded and expanded mouse neuroblastoma, Neuro-2a (N2a) cells, and observed clear staining of various actin structures post-expansion **(Fig. S2B, S2C, S2F)**. N2a cells also displayed isotropic expansion with the nucleus area increasing by an average expansion factor of 3.88 and 3.24 for β- and γ-IntAct expressing N2a cells **(Fig. S2B-E)**. These results highlight the significant advantage provided by IntAct-U-ExM to study mammalian actin networks at high 3D-molecular resolution and highlight the strong potential for future studies of actin and its binding-proteins with multiplexed imaging^63^.

**Figure 2.**
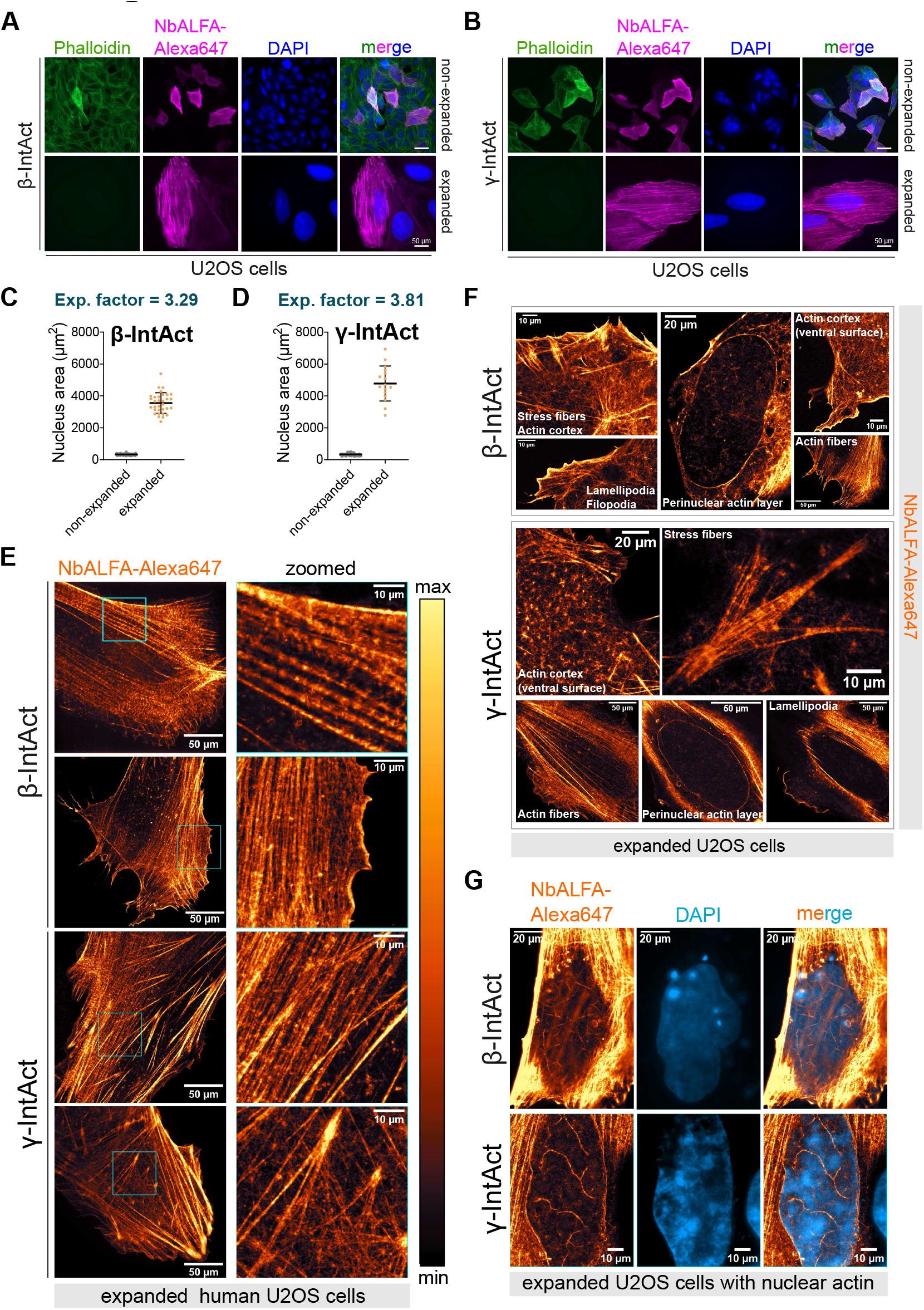
IntAct-U-ExM can be used to study mammalian actin networks in an isoform-specific manner. **(A)** Representative maximum intensity projected images of non-expanded and expanded human U2OS cells expressing β**-**IntAct stained as indicated. **(B)** Representative maximum intensity projected images of non-expanded and expanded human U2OS cells expressing γ**-**IntAct stained as indicated. **(C)** Plots representing measurements of nucleus area in non-expanded and expanded U2OS cells expressing β**-**IntAct along with calculated expansion factors. **(D)** Plots representing measurements of nucleus area in non-expanded and expanded U2OS cells expressing γ**-**IntAct along with calculated expansion factors. **(E)** Representative maximum intensity projected images of expanded human U2OS cells expressing either β- or γ-IntAct stained with NbALFA-Alexa647 showing various actin structures. **(F)** Representative singe plane or maximum intensity projected images of expanded human U2OS cells expressing either β- or γ-IntAct stained with NbALFA-Alexa647 showing diversity of actin filament networks as indicated. **(G)** Representative singe plane or maximum intensity projected images of expanded human U2OS cells expressing either β- or γ-IntAct stained with NbALFA-Alexa647 showing presence of actin filaments inside the nuclear volume.

Interestingly, we detected both β- and γ-IntAct signal appearing as filaments in the nucleus of U2OS cells **(Fig. 2G)** and as a continuous layer surrounding the nucleus in expanded U2OS cells **(Fig. 2F, 2G)**. Actin filaments inside the nucleus have been observed previously with the use of nuclear-targeted actin chromobody (nAC)^64^ and remain challenging to visualize by other conventional labelling approaches^65^. Our results, thus, demonstrate an alternate way to study these nuclear actin filaments with isoform specificity. These observations demonstrate the power of IntAct-U-ExM in elucidating lesser studied actin filament populations in diverse cellular compartments, promising to reveal new functional aspects of actin.

Taken together, our results strongly demonstrate the versatile application of IntAct as a potent tool to study diverse actin-based structures with super-resolution expansion microscopy in yeast and mammalian cells. It also provides a robust modality to study specific isoforms of actin and their roles in diverse cellular compartments. The versatile nature of the internal ALFA tag and the small size of the NbALFA provide a useful system for super-resolution microscopy of the actin cytoskeleton across species. When coupled with rapid technological advancements in ExM probes/workflows which continue to increase molecular resolution and combine ExM with other modalities like STED, SMLM, SIM, etc., IntAct-U-ExM provides an ideal and compatible platform which could reveal many new fundamental insights into the nanoscale architecture of diverse actin isoforms and networks across model systems.

## Acknowledgements

The authors thank Dr. Koen van den Dries for sharing the plasmids for mammalian expression and critical reading of the manuscript. We also thank Dr. Hashim Reza for his immense and constant help with optimization and troubleshooting of U-ExM workflow for the study. AD acknowledges GATE fellowship from IISc. SM and SD acknowledge Research fellowship from the Department of Biotechnology, Govt of India.

## Data Availability

All supporting data is available upon request from the authors.

## Materials and Methods

### Plasmids and Yeast strains used in the study

All plasmids and yeast strains used in this study are described in Supplemental Table 1 and Supplemental Table 2.

### Yeast growth

*S. cerevisiae* and *S. pombe* strains were grown overnight in Synthetic Complete media lacking uracil (SC-ura) at 25°C with shaking at 250 rpm. The overnight culture was diluted to a O.D._600_ = 0.2, grown till mid-log phase and harvested for immunofluorescence or expansion microscopy.

### Cell culture and Transfection

#### U2OS cells

U2OS (ATCC® HTB-96™) cells were cultured under standard conditions at 37°C in a humidified 5% CO_2_, incubator in Dulbecco’s Modified Eagle’s Medium (DMEM, Gibco 11965118) containing 10% FBS (Gibco A5256701), 100 U/mL Penicillin, 0.1 mg/mL Streptomycin (Gibco 15140-122) and 2 mM L-glutamine (HiMedia, TCL012) in a humidified incubator. For transfections, 4 x104 cells/500 µl were plated on 12-mm glass coverslips in a 24-well plate, cultured overnight and then, transfected with 1 *µ*g of pcDNA3.1-β-IntAct or pcDNA3.1-γ-IntAct with jetPEI® (PolyPlus) transfection medium following the manufacturer’s instructions. 24 h post transfection, the culture media was aspirated, and the cells were fixed with 4% paraformaldehyde in phosphate buffered saline, pH 7.2 (PBS) for 15 min at room temperature (RT) followed by washing with PBS. The fixed samples were stored at 4 °C till further use.

#### N2a cells

Neuro-2a cell line, a mouse neuroblastoma cell line (ATCC® CCL-131TM, RRID: CVCL_0470) was cultured at 37 °C and 5% CO2 in Dulbecco modified Eagle medium (Gibco 12430-054) with 100 U Penicillin/mL and 0.1 mg Streptomycin/mL (Pen Strep Gibco 15140-122), 10% Fetal Bovine Serum (FBS; HIMEDIA RM10434), and 1 % Glutamax (Gibco 35050/061). For transfection, 6 × 10^4^ cells/well were seeded on 18-mm coverslips in a 12-well plate and grown overnight in complete DMEM after which, the media was changed to pre-filtered DMEM without antibiotics. the cells were transfected with 20 ng of either β-IntAct or γ-IntAct using Lipofectamine™ 3000 Transfection Reagent (#L3000075, Thermo Scientific) where the plasmid DNA was diluted in 50 µl of Opti MEM Reduced Serum Medium (#31985070, Thermo Scientific) along with 1 µl of P3000 reagent. Separately, 2 µl of Lipofectamine 3000 reagent was diluted in 50 µl of Opti MEM. The two mixtures were then combined, gently mixed, and incubated at RT for 20 min followed by addition to the cells after which the plate was gentle rocked to ensure even distribution. 24 h - 48 h post transfection, the cells were fixed with 4% PFA in PBS for 15 min at RT, washed with PBS and stored at 4°C till required for expansion.

## Ultrastructure expansion microscopy (U-ExM) workflow for yeast

### Reagents required

- Acrylamide (AA). Stock: 40% (Sigma-Aldrich A4058). Stored at 4°C.
- Formaldehyde (FA). Stock: 36.5-38% (Sigma-Aldrich F8775). Stored in the fume hood.
- N, N’-methylenebisacrylamide (BIS). Stock: 2% (Sigma-Aldrich M1533). Stored at 4°C.
- Poly-L-lysine. (Sigma-Aldrich A-003-E). Stored at 4°C.
- Nuclease-free water.
- Sodium acrylate (SA). Stock: 97-99% (Sigma-Aldrich 408220)
- Ammonium persulfate (APS)
- Tetramethylethylenediamine (TEMED)

### Day 1: Sample preparation and Anchoring

A modified version of a previously described U-ExM protocol for yeast was followed (ref). Yeast cells grown till mid-log phase were fixed with either (i) 4% formaldehyde (FA) or (ii) 4% formaldehyde + 0.1% glutaraldehyde for 15 minutes in PEM buffer (100 mM PIPES, 1 mM EGTA, 1 mM MgSO4, pH adjusted to 6.9). Cells were then washed with 1x PBS three times and then incubated with 100mL of PEM-S (1.2 M Sorbitol in PEM) buffer containing 15mL of Long-Life Zymolase (#786–036, GBioSciences) at 37°C for 30-45 minutes. The cells were then washed twice with PEM-S buffer, resuspended in freshly made 100mL AA-FA mixture (1%AA + 0.7%FA) made in 1x PBS, and incubated overnight at 37°C, 350 rpm.

### Day 2: Seeding, cross-linking to gel, denaturation, and staining

The overnight incubated mixture of cells was seeded on a 12mm round glass coverslip coated with poly-L-lysine and allowed to sit for 20 minutes at room temperature. The excess cell mixture was removed and stored at 4°C. Simultaneously, 36mL of monomer solution (MS; 19% (wt/wt) SA, 10% (wt/wt) AA, 0.1% (wt/wt) BIS in PBS) was mixed with 2mL of 10% TEMED and 2mL of 10% APS. The mixture was immediately poured as a droplet of 36mL on a parafilm attached to a flat metal block kept on ice. The coverslip with seeded cells was gently placed on top of the droplet such as to cover the whole droplet and allowed to sit on the cold metal block for 5 minutes. The metal block was then transferred to a sealed humid chamber and kept at 37°C for 45 minutes to allow the polymerization of the gel. The gels were then separated from the coverslips and incubated in denaturation buffer (50 mM Tris pH 9.0, 200 mM NaCl, 200 mM SDS, Adjust pH to 9.0 with HCl) for 90 minutes at 95°C. This was followed by washing with PBS thrice for five minutes at room temperature. Furthermore, the gels were then stained with FluoTag-X2 anti-ALFA-Alexa647 (#N1502, Nanotag Biotechnologies, 1:100 dilution) overnight in 3% BSA in 1x PBS-T at 4°C.

### Day 3: Imaging

The gels were stained with DAPI (1:200) for 30 minutes at room temperature, followed by three washes with 1x PBS-T. Subsequently, the gels were expanded in ddH_2_O thrice for 30 minutes each and then mounted on a 35mm glass bottom dish (Ibidi, Cat.No: 81151) coated with poly-L-lysine. Imaging was done with an Olympus SpinSR spinning disk confocal microscope using a 60x oil-immersion objective (N.A.= 1.42). The samples were excited with a solid-state laser of wavelength 640 nm and a LED light source of wavelength 405nm. The images were acquired with a Prime BSI scMOS camera and deconvolved using Olympus CellSens Dimension software.

## Immunofluorescence imaging of non-expanded yeast cells

Immunofluorescence was performed as described previously (ref). Briefly, yeast strains were grown overnight at 25°C in YPD broth/EMM-Ura. The overnight culture was diluted and allowed to grow until mid-log phase. Cells were fixed with 4% formaldehyde for 60 min at 25°C, washed twice with 1x PBS, and finally resuspended in 200 µl of 1.2M Sorbitol Phosphate-Citrate (SPC) buffer (1.2 M Sorbitol, 1 M K2HPO4, 1 M Citric acid); 25 µl of Long-Life Zymolase (#786–036, GBiosciences) was added to digest the yeast cell wall and the suspension was incubated with mild shaking at 37°C for 60 min. The cells were then washed twice with ice-cold SPC buffer and incubated with 500 µl of blocking buffer 2% bovine serum albumin (BSA) + 0.1% Triton X-100 in PBS at room temperature for 15 min with shaking. The cells were pelleted and resuspended in 500 µl of Antibody Dilution Buffer (1% BSA + 0.05% Triton X-100 in PBS) containing FluoTag-X2 anti-ALFA-Alexa647 (#N1502, Nanotag Biotechnologies) at a final dilution of 1:500. The cell suspension was then incubated overnight with rotation at 4°C. Next day, cells were washed twice with 1xPBS and finally resuspended in 20 µl of 1x PBS, and 5 µl of the final cell suspension was mounted on a glass-bottom dish coated by poly-L-lysine (#P4707, Sigma Aldrich). Phalloidin staining of yeast actin structures was done as per previously described protocols [63,64]. Briefly, cells were grown at 25°C till mid-log phase and fixed with 4% paraformaldehyde. The cells were washed thrice with 1x PBS and labeled phalloidin was added to a final concentration of 0.4 *µ*M (in 50 µl of 1x PBS) and the tubes were kept in a rotating shaker overnight at 4°C. The cells were washed twice with 1x PBS on the next day and seeded on a concanavalin A coated glass-bottom dish. Imaging was done with an Olympus SpinSR spinning disk confocal microscope using a 100x oil-immersion objective (N.A.=). The samples were excited with solid-state lasers of wavelength 640 nm and 488 nm and a LED light source of wavelength 405nm. The images were acquired with a Prime BSI scMOS camera and deconvolved using Olympus CellSens Dimension software.

## Ultrastructure expansion microscopy (U-ExM) workflow for U2OS cells

### Materials required

- Nuclease-free water (NFW, #AM9937, Ambion-ThermoFisher)
- Poly-D-Lysine (#A3890401, Gibco)
- Ammonium persulfate (APS, #17874, ThermoFisher)
- Formaldehyde (FA, #F8775, SIGMA)
- Tetramethylethylenediamine (TEMED, #17919, ThermoFisher)
- Acrylamide (AA, 40%, #A4058, SIGMA) – Ready to use – Keep at 4°C
- N, N’-methylenebisacrylamide (BIS, 2%, #M1533, SIGMA)
- Sodium Acrylate (SA, #408220, SIGMA)
- Glass-bottom Confocal dish (#BDD-002-035, BioFil)

### Day 1: Fixing and first round of expansion

The β-IntAct or γ-IntAct transfected U2OS cells, fixed with 4% PFA were first incubated with 2.8% formaldehyde / 5% acrylamide solution for 5 h at 37°C. Then, the solution is removed, and the cells were incubated with 35 µl of monomer solution (composed of 25% acrylamide, 5% bis-acrylamide, 19% sodium acrylate, 0.5% TEMED and 0.5% ammonium persulfate) for 5 min on ice, followed by 1 h at 37°C to allow gelation. Post gelation, the coverslips with the gels were transferred to a 6-well plate and incubated with 1 mL of denaturation buffer (50 mM Tris/HCl, pH 9.0 containing 200 mM SDS and 200 mM NaCl) for 15 min at RT with shaking. Once the gel detaches from the coverslip, carefully transfer it with a spatula into a fresh 1.5mL vial filled with 1-2 mL of fresh denaturation buffer and incubated for 90 min at 95°C. Post denaturation, the gel was carefully transferred to a glass petri-dish with distilled water for 30 min followed by changing the water and incubating the gels overnight for complete expansion.

### Day 2: Staining, expansion and imaging

The expanded gels were exchanged with PBS for water twice for 15 min each to shrink the gel for immunostaining. Then a small piece of the gel was cut using a scalpel and carefully transferred to a 24-well plate and incubated with Phalloidin-Alexa488 (#49409, Sigma Aldrich) at a final dilution of 1:50 and FluoTag-X2 anti-ALFA-Alexa647 (#N1502, Nanotag Biotechnologies) at a final dilution of 1:500, both diluted in PBS containing 2% BSA for 2 h 30 min at 37°C. The gels were subsequently washed 3 times with PBS containing 0.1% Tween 20 (PBST) for 10 min at RT with agitation. Then, the gel was incubated with DAPI (#D1306, Thermo Scientific) at a final dilution of 1:200 in PBS for 20 min RT with agitation following which, the gel piece was transferred to a glass petri-dish containing distilled water for 30 min for the final round of expansion before proceeding to image the gels. Once expanded, a small piece of the expanded gel was decanted of any excess water by gently tapping it with tissue paper. and transferred to a poly-D-lysine coated 35 mm glass-bottom confocal dish and gently pressed it to ensure it is nicely fixed onto the dish. The piece was then imaged using a 63x oil immersion objective (NA=1.42) on the Zeiss Axio Observer 7 epifluorescence microscope equipped with the Orca-Flash4.0 V3 sCMOS camera (Hamamatsu) using the ZEN 3.8 (Carl Zeiss) software or Olympus SpinSR spinning disk confocal microscope using a 60x oil-immersion objective (N.A.= 1.42) using the Prime BSI sCMOS camera and CellSens (Olympus) software. The images were deconvolved using the deconvolution modules within the respective software when required.

## Immunofluorescence imaging of non-expanded U2OS and N2a cells

The β-IntAct or γ-IntAct transfected U2OS cells, fixed with 4% PFA were incubated with Phalloidin-Alexa488 (1:200 dilution), FluoTag-X2 anti-ALFA-Alexa647 (1:500 dilution) and DAPI (1:1000 dilution) all diluted in PBS containing 2% BSA for 2h at RT. The coverslips were washed 3 times with PBST for 3 min at RT and mounted onto a glass slide using ProLong Gold (#P36934, Thermo Scientific) and allowed to polymerize overnight at RT. The cells were imaged using a 63x oil-immersion objective (NA=1.42) on the Zeiss Axio Observer 7 epifluorescence microscope equipped with the Orca-Flash4.0 V3 sCMOS camera (Hamamatsu) using the ZEN 3.8 (Carl Zeiss) software

## Ultrastructure expansion microscopy (U-ExM) workflow for N2a cells

Neuro-2a cells were processed for U-ExM using the same protocol as mentioned above for yeast.

## Image Analysis

All images were analyzed in Fiji (ImageJ). The nuclei were segmented using automatic segmentation using the Labkit plug-in. The cell dimensions were measured manually in ImageJ. Expansion factors were calculated by comparing measured parameters (nucleus area, cell area, cell length and width) between non-expanded and expanded cells. For FWHM analysis, line plots perpendicular to the yeast actin cables were generated with ImageJ and non-linear gaussian curve fitting was done in GraphPad Prism (v10.2) using the equation:

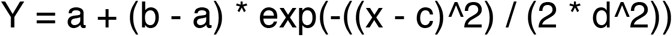

where amplitude = (b-a); mean = c; Standard deviation (SD) = d; Baseline = a (Y at x=0). The FWHM was calculated as (2.355 * SD) and the values were plotted using GraphPad Prism (v10.2).

## Figure Legends

**Supplementary Figure 1.**
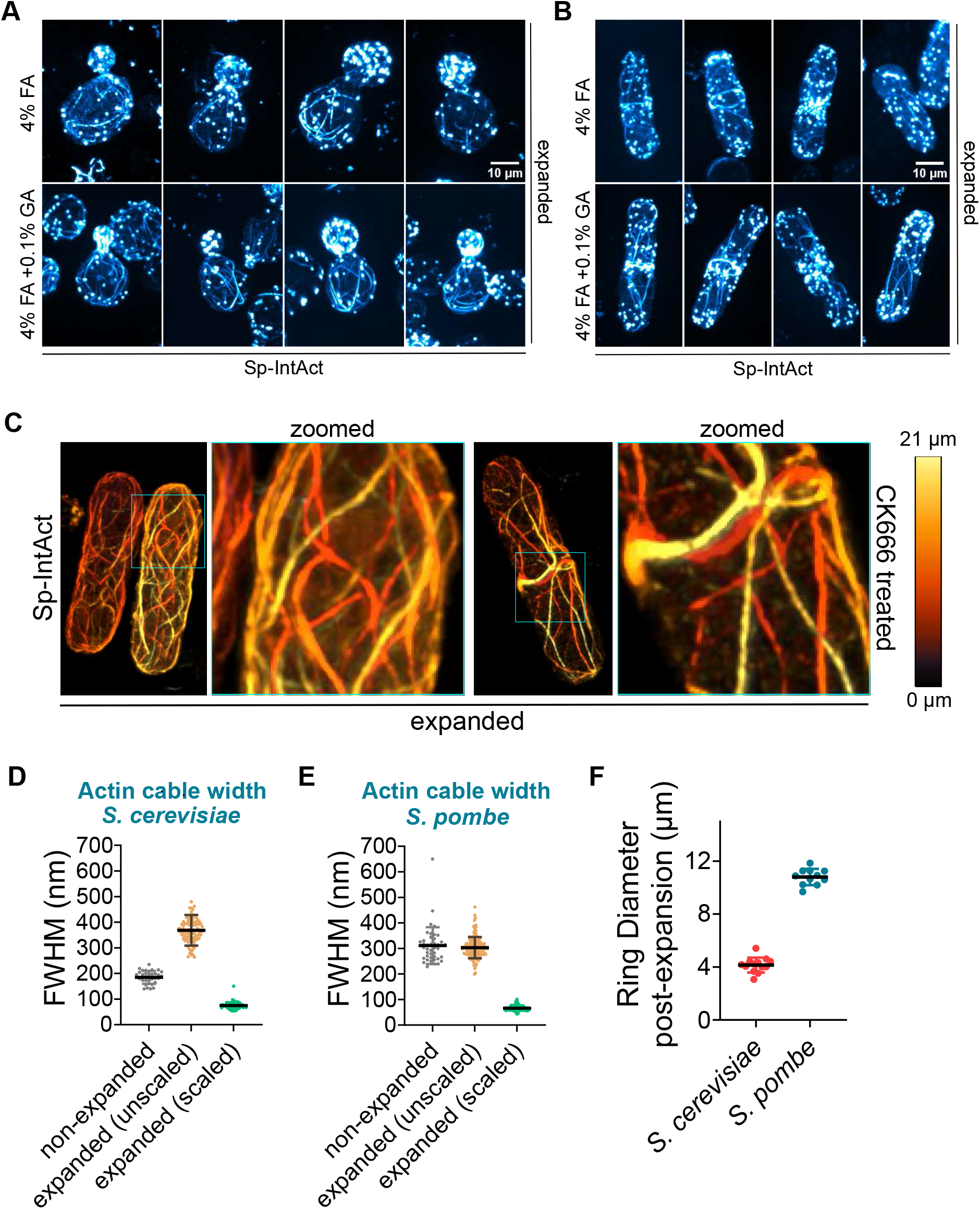
(A) Representative maximum intensity projected images showing comparison of expanded *S. cerevisiae* cells fixed with either 4% FA (top row) or 4% FA + 0.1% GA (bottom row), stained with NbALFA-Alexa647. (B) Representative maximum intensity projected images showing comparison of expanded *S. pombe* cells fixed with either 4% FA (top row) or 4% FA + 0.1% GA (bottom row), stained with NbALFA-Alexa647. (C) Representative maximum intensity projected images of expanded *S. pombe* cells treated with Arp2/3 inhibitor (100μM) for 15 mins prior to U-ExM and stained with NbALFA-Alexa647. (D, E) Plots depicting measurements of Full Width at Half Maxima (FWHM) for actin cables in non-expanded (phalloidin stained) and expanded (NbALFA-Alexa647 stained) *S. cerevisiae* (D) and *S. pombe* (E); values from expanded samples were scaled down with the average expansion factor of 4.92. (F) Plot depicting measured actomyosin ring diameter in expanded *S. cerevisiae* and *S. pombe* cells.

**Supplementary Figure 2.**
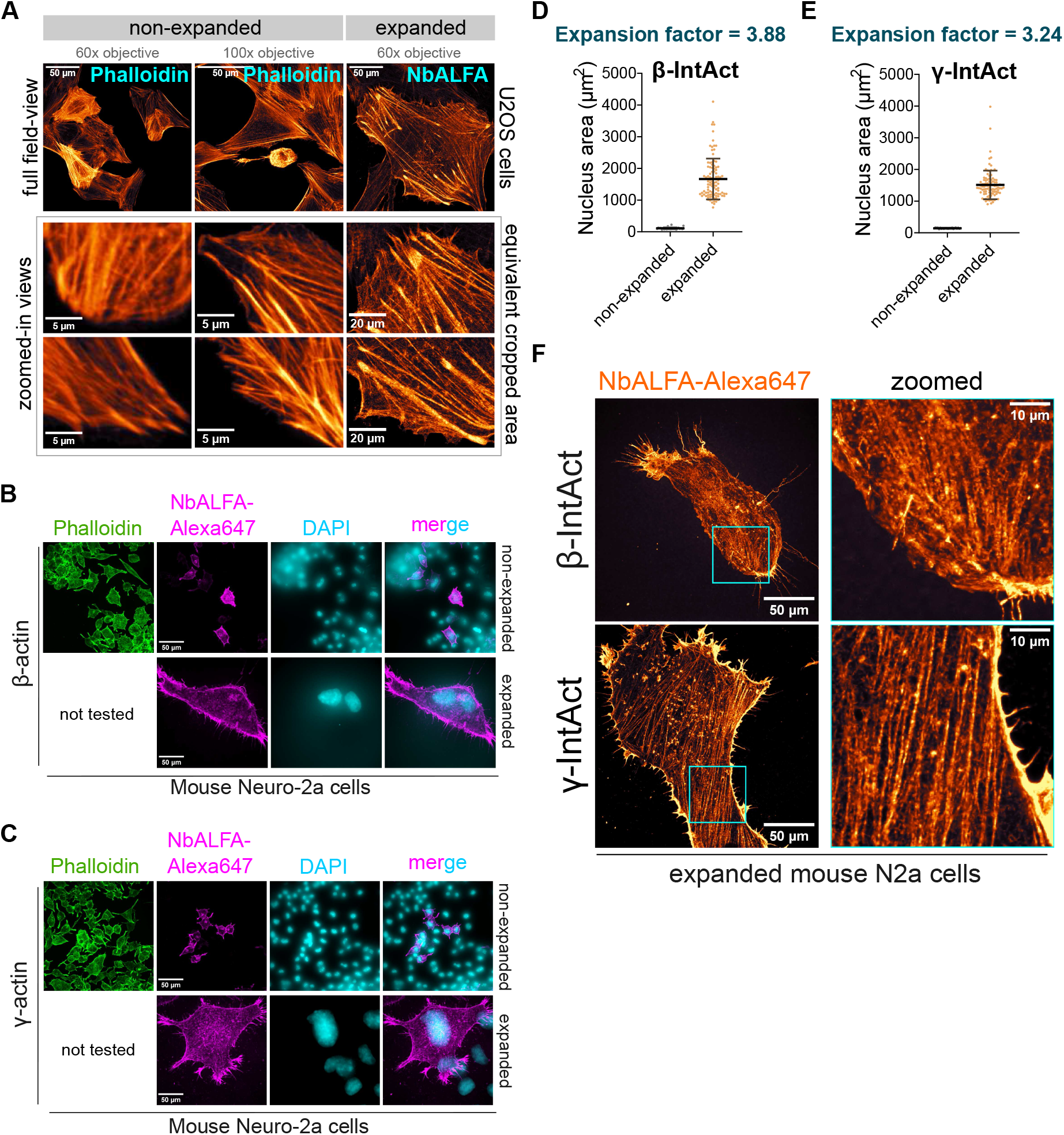
(A) Representative maximum intensity projected images qualitatively showing difference in resolution across non-expanded and expanded human U2OS cells expressing IntAct and stained as indicated. (B) Representative maximum intensity projected images of non-expanded and expanded mouse Neuro-2a (N2a) cells expressing β**-**IntAct stained as indicated. (C) Representative maximum intensity projected images of non-expanded and expanded mouse Neuro-2a (N2a) cells expressing γ**-**IntAct stained as indicated. (D) Plots representing measurements of nucleus area in non-expanded and expanded mouse N2a cells expressing β**-**IntAct along with calculated expansion factors. (E) Plots representing measurements of nucleus area in non-expanded and expanded mouse N2a cells expressing γ**-**IntAct along with calculated expansion factors. (F) Representative maximum intensity projected images of expanded human mouse N2a cells expressing either β- or γ-IntAct stained with NbALFA-Alexa647 showing various actin structures.

## Supplementary Movie Legends

**Movie S1**. Movie representing 3D volume (z-stacks) of expanded *S. cerevisiae* and *S. pombe* cells stained for IntAct with NbALFA-Alexa647 (blue).

**Movie S2**. Movie representing 3D volume (z-stacks) of expanded *S. pombe* cells treated with Apr2/3 inhibitor CK666 stained for IntAct with NbALFA-Alexa647 (orange). Different colors represent different z-depth

**Movie S3**. Movie representing 3D volume (z-stacks) of expanded U2OS cells stained for

β- or γ-IntAct with NbALFA-Alexa647 (orange).

**Supplementary Table 1.**
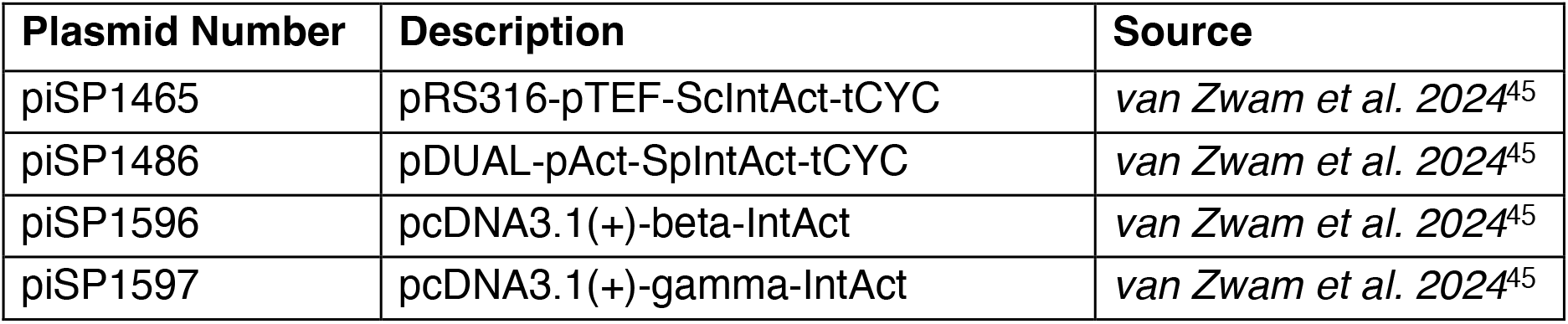
List of plasmids used in this study.

**Supplementary Table 2.** List of yeast strains used in this study.

| **Strain Number** | **Genotype** | **Source** |
| --- | --- | --- |
| YSP641 | *MATa leu2Δ1 trp1Δ63 his3Δ200 ura3-52 :: pRS316-pTEF-Sc-IntAct-tCYC-ura3+* | *van Zwam et al. 2024<sup>45</sup>* |
| YSP1049 | *ura4-D18 leu1-32 h- :: pDUAL-pAct-Sp-IntAct-Tadh1-ura4+* | *van Zwam et al. 2024<sup>45</sup>* |

