## Supplementary figures and images for "IntAct-U-ExM: Ultrastructure Expansion microscopy of actin networks via an internally-tagged actin"

### Supplemental_figures_S1 and S2

#### Supplementary Figure 1

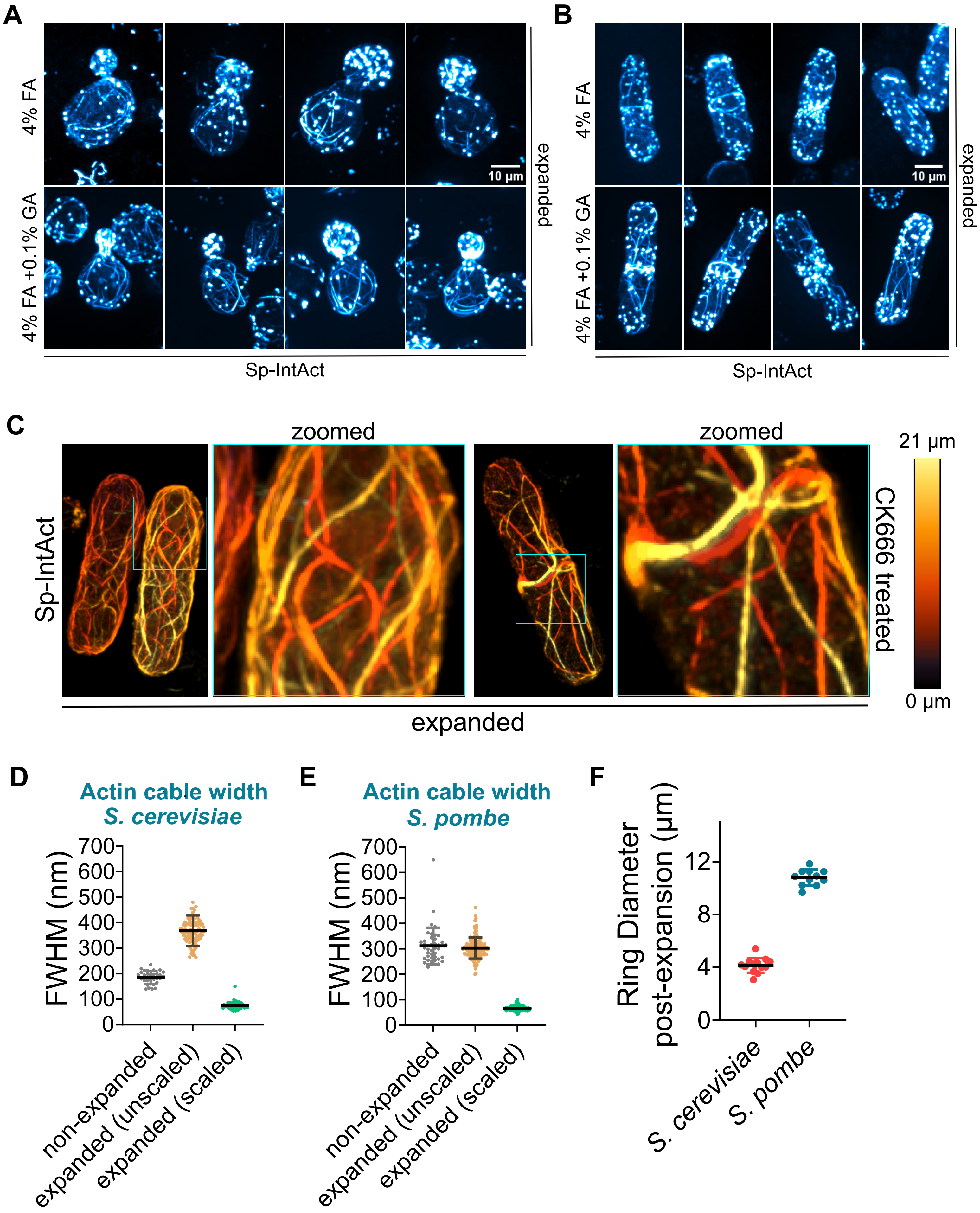

#### Supplementary Figure 2

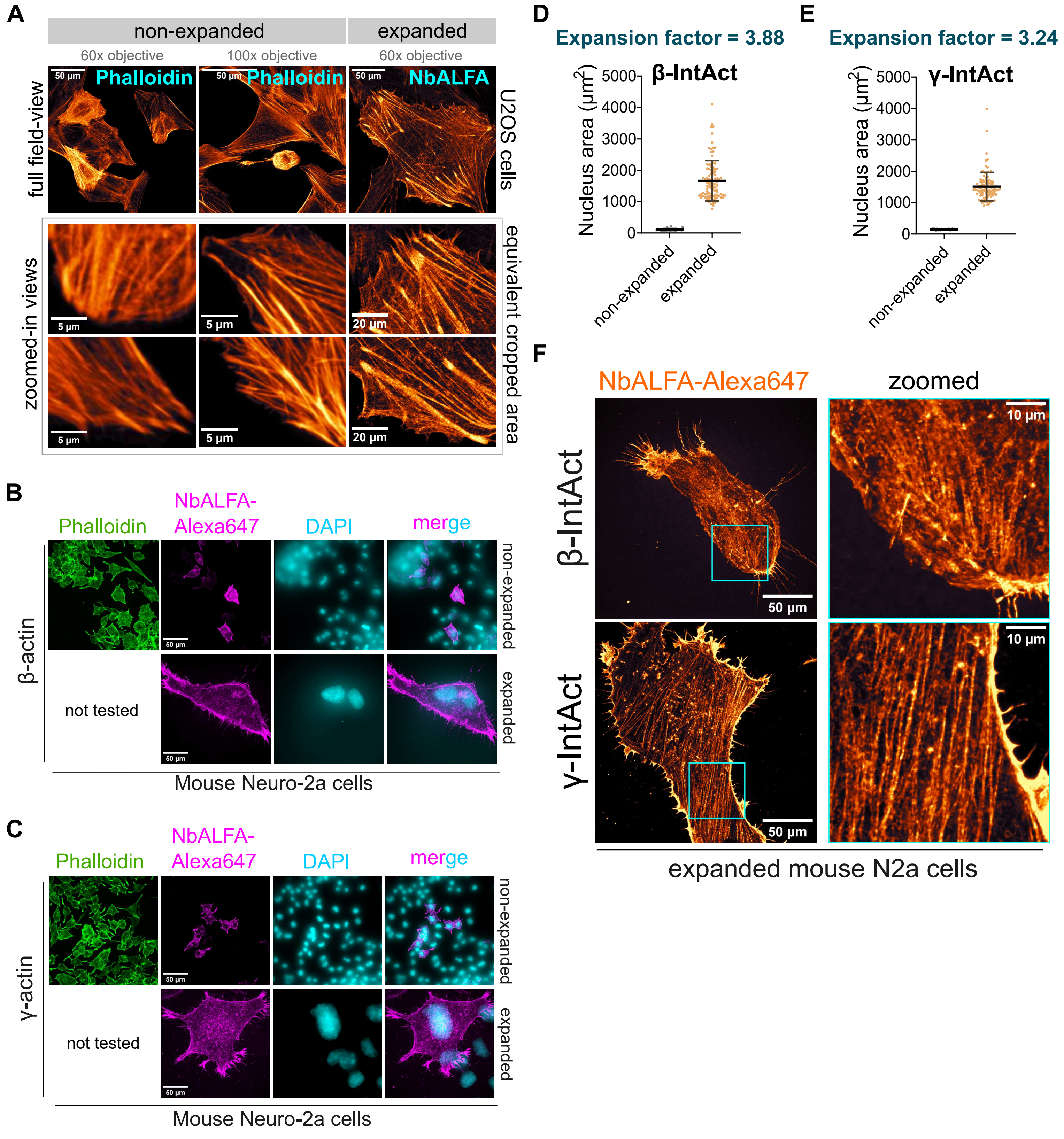
